# Sun navigation requires compass neurons in *Drosophila*

**DOI:** 10.1101/315176

**Authors:** Ysabel Milton Giraldo, Katherine J. Leitch, Ivo K. Ros, Timothy L. Warren, Peter T. Weir, Michael H. Dickinson

## Abstract

To follow a straight course, animals must maintain a constant heading relative to a fixed, distant landmark, a strategy termed menotaxis. In experiments using a flight simulator, we found that *Drosophila* adopt arbitrary headings with respect to a simulated sun, and individuals remember their heading preference between successive flights—even over gaps lasting several hours. Imaging experiments revealed that a class of neurons within the central complex, which have been previously shown to act as an internal compass, track the azimuthal motion of a sun stimulus. When these neurons are silenced, flies no longer adopt and maintain arbitrary headings, but instead exhibit frontal phototaxis. Thus, without the compass system, flies lose the ability to execute menotaxis and revert to a simpler, reflexive behavior.

**One sentence summary:** Silencing the compass neurons in the central complex of *Drosophila* eliminates sun navigation but leaves phototaxis intact.

---

Despite their small brains, insects can navigate over long distances – in some cases, thousands of kilometers – by orienting to sensory cues such as visual landmarks (*1*), skylight polarization (*2–9*) and the position of the sun (*3, 4, 6, 10*). Although *Drosophila* are not generally renowned for their navigational abilities, mark-and-recapture experiments in Death Valley revealed that they can fly nearly 15 km across open desert over the course of a single evening (*11*). To accomplish such feats on available energy reserves (*12*), flies would have to maintain relatively straight headings and rely on celestial cues to do so (*13*).

Celestial cues such as sun position and polarized light are thought to be integrated in the central complex, a set of highly conserved unpaired neuropils in the central brain of arthropods (*14*). Central complex neurons in locusts (*15*), dung beetles (*4*), and monarch butterflies (*16*) respond to the angle of polarized light and the position of small bright objects mimicking the sun or moon. Extracellular recordings from the central complex in cockroaches revealed neurons that act as head-direction cells in the absence of visual cues or relative to a visual landmark (*17*). Recently, a group of cells (E-PG neurons) in the *Drosophila* central complex have been shown to function as an internal compass (*18–20*), similar to head-direction cells in mammals (*21*). Using the wide array of genetic tools available to measure and manipulate cell function in *Drosophila*, we set out to test whether flies can navigate using the sun and to identify the role of E-PG cells in this behavior.

We tested the hypothesis that *Drosophila* can use the sun to navigate by placing tethered wild-type female flies in a flight simulator and presenting an ersatz sun stimulus (Fig.1A). The fly was surrounded by an array of LEDs on which we presented either a single 2.3° bright spot on a dark background or a 15°-wide dark vertical stripe on a bright background. Given previous studies on other species (*4, 15, 16*), we expect that flies react to our small bright spot as they would to the actual sun, and thus we call it a ‘sun stimulus’. Experiments were conducted in closed loop, such that the difference in stroke amplitude between the fly’s two wings determined the angular velocity of the stimulus (*12*). Flies generally maintained the dark stripe in front of them (Fig. 1C, D), a well-characterized behavior termed stripe fixation (*22–24*). However, when presented with the sun stimulus, individual flies adopted arbitrary headings, thus exhibiting menotaxis (Fig. 1B, D). We quantified how well flies maintained a heading by calculating vector strength, which is the magnitude of the mean of all instantaneous unit heading vectors for the entire flight. A vector strength of 1 would indicate that a fly held the stimulus at the exact same heading during the entire flight bout. Because we tested each individual with both a stripe and sun stimulus, we could compare the flies’ performance under the two conditions. We found no correlation between the mean heading exhibited by individual flies during sun menotaxis and stripe fixation (Fig. 1E), suggesting that heading preference for the sun stimulus is independent of the response to a vertical stripe. To ensure that flies’ stabilization of the sun stimulus was not an artifact of our feedback system, we also conducted control closed-loop experiments in which the bright spot was switched off. As expected, the flies exhibited no orientation behavior under this condition, with all vector strength values lower than 0.16 (Fig. 1D). Collectively, these experiments indicate that flies are capable of orienting to a small bright spot and that this behavior is distinct from stripe fixation. *Drosophila* can also perform menotaxis using the axis of linearly polarized light (*8, 9, 25*). It is not known whether the orientation responses of flies to the sun and polarized light are independent, as they are in dung beetles (*4*), or linked to create a matched filter of the sky, as they are in locusts (*15*).

**Fig. 1.**
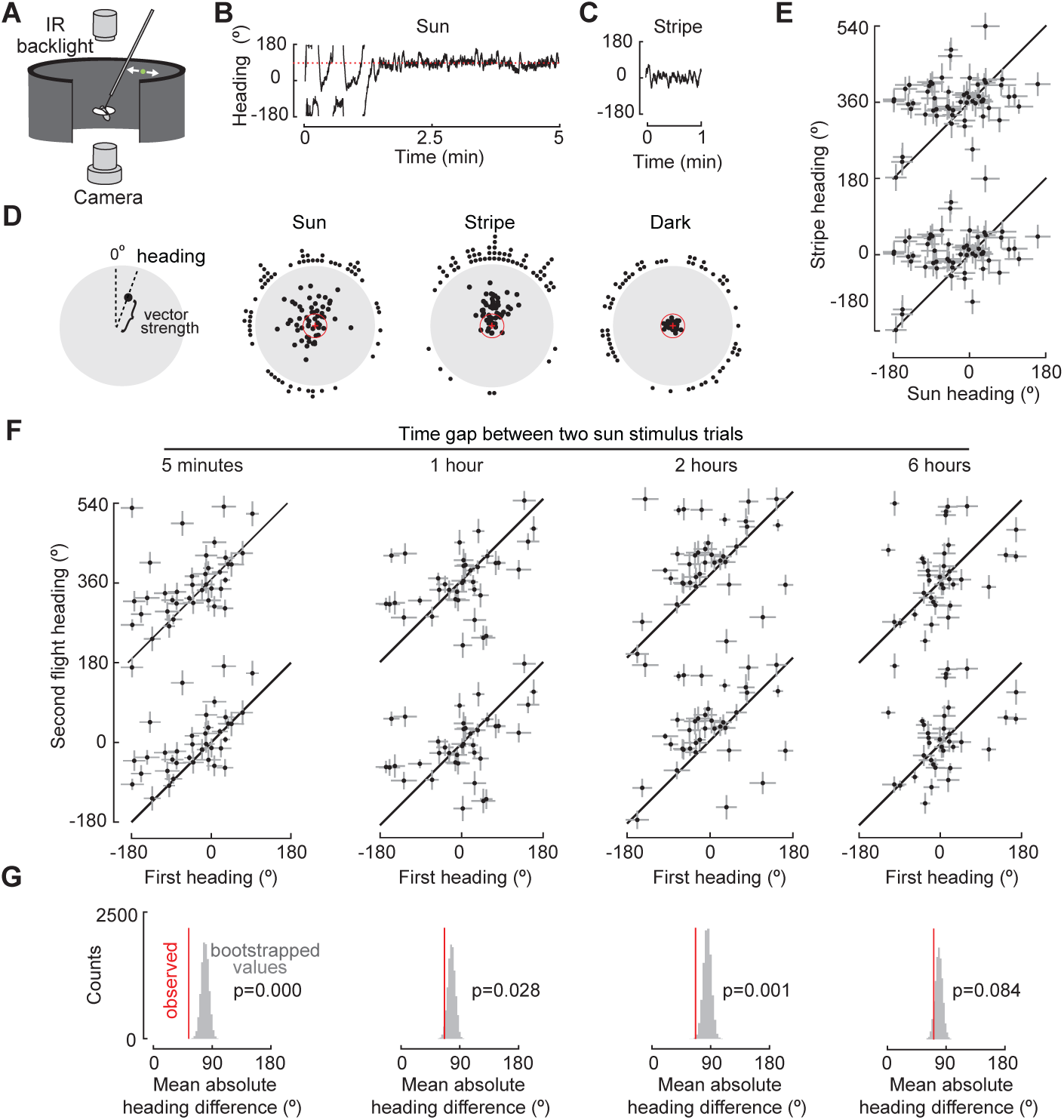
Flies navigate using a sun stimulus and retain memory of their heading. (A) Tethered fly, backlit with infrared light and surrounded by a cylindrical LED display; a single 1.43° spot simulates the sun. (B) Example trace showing closed-loop behavior. After 88 seconds, the fly stabilized the sun stimulus at a heading of 92°. (C) Heading during a stripe presentation. (D) Polar representation of data. Angular position indicates mean heading; radial distance indicates vector strength. Headings for flies presented with a sun stimulus, a stripe, and in the dark. A histogram of mean heading is plotted around each circle. Dashed red circle, vector strength of 0.2. (E) Sun versus stripe heading. Data are repeated on the vertical axis to indicate their circular nature. Diagonal line indicates identical heading over both trials. Error bars, circular variance times 36 for visibility. (F) Heading in first trial plotted against second trial heading for increasing inter-trial intervals; plotting conventions as in 1E. (G) Distribution of 10,000 bootstrapped heading differences between random pairings of first and second trials from 1F. Red line, mean heading difference of observed data; p-value, proportion of resampled differences that are smaller than the observed mean heading difference.

Given that individual flies adopted arbitrary headings with respect to the sun stimulus, we tested whether they retained a memory of their orientation preference from one flight to the next. We presented flies with the sun stimulus in closed loop, interrupted flight for a defined interval (5 min, 1, 2, or 6 hours), and then again presented the sun stimulus. To provide an independent metric of flight performance, we also presented a stripe under closed loop conditions for 1 min before the first sun bout and after the second. Across inter-flight intervals of 5 minutes, 1 hours, and 2 hours, flies remained loyal to their first heading during the second flight (Fig. 1F). If each fly adopted the identical heading in both flights, the mean heading difference would be zero, whereas if there was no correlation in heading from one flight to the next, the mean absolute value of the heading difference would be 90°, provided that the orientations were uniformly distributed. To test whether the consistency in flight-to-flight orientation could arise from chance, we bootstrapped 10,000 random pairs of mean heading values from the first and second flights and compared the resulting distribution with the mean absolute heading difference of the actual data (Figure 1G). In all cases, the measured mean difference was less than that of the bootstrapped values (5 min: 54.2° vs. 79.2°; 1 hour: 66.6° vs. 77.4°; 2 hours: 66.8° vs. 84.8°; 6 hours: 71.0° vs. 75.5°). We calculated probability values directly from the proportion of the 10,000 bootstrapped simulations that resulted in a smaller mean absolute angle difference than the observed data (Fig.1G). With the exception of the 6-hour gap, this probability was quite low (5 min: p=0.00; 1 hour: p=0.03; 2 hours: p=0.001; 6 hours, p=0.084). Collectively, these results suggest that headings are not selected at random with each subsequent takeoff, but rather that flies remember their headings from previous flights, at least for up to 2 hours. A similar result was found for the orientation responses to linearly polarized light (*8*). Fully determining the mechanisms by which flies attain their initial heading preference (i.e. genetic vs. developmental vs. learning) require experiments that are beyond the scope of this current study.

The finding that flies remember their flight heading for at least 2 hours makes ethological sense. *Drosophila* are crepuscular, exhibiting dawn and dusk activity peaks (*26*). Assuming our laboratory measurements are representative of dispersal events, a memory that allows an individual to fly straight for a few hours would be sufficient to bias a day’s migration in one direction. To our knowledge, there is no evidence that *Drosophila* make multi-day, long-distance migrations that would require the ability to maintain a constant course from one day to the next or a time-compensated sun compass. The most parsimonious ecological interpretation of their sun orientation behavior is that it allows flies to disperse opportunistically to new sources of food and oviposition sites within a single day.

The visual information conveying sun position likely provides inputs to the recently identified neurons constituting the fly’s internal compass (*18–20*). These columnar neurons receive input in the ellipsoid body and send divergent output to the protocerebral bridge and gall, and are hence named E-PG neurons (*27*). These neurons track azimuthal position of vertical stripes and more complex visual stimuli, and in the absence of visual input can continue to track azimuthal orientation by integrating estimates of angular velocity (*18, 20, 28*). Given these functional attributes, an obvious question is whether E-PG neurons respond to a sun stimulus and whether they exhibit different responses to other visual stimuli. We used the split-GAL4 line SS00096 (*28*), which expresses in the E-PG neurons, to drive the genetically encoded calcium indicator GCaMP6f, and measured activity in tethered, flying flies using a 2-photon microscope (Fig. 2A). As described previously, the set of 16 E-PG neurons tile the toroidally shaped ellipsoid body. Notably, a region of activity, or ‘bump’, rotates around the ellipsoid body corresponding to azimuthal position (*18*, Movies S1, S2). Instead of recording from the ellipsoid body, we imaged the activity at E-PG terminals in the protocerebral bridge (Fig. 2B) because fluorescence signals were stronger in these more superficial glomeruli.

**Fig. 2.**
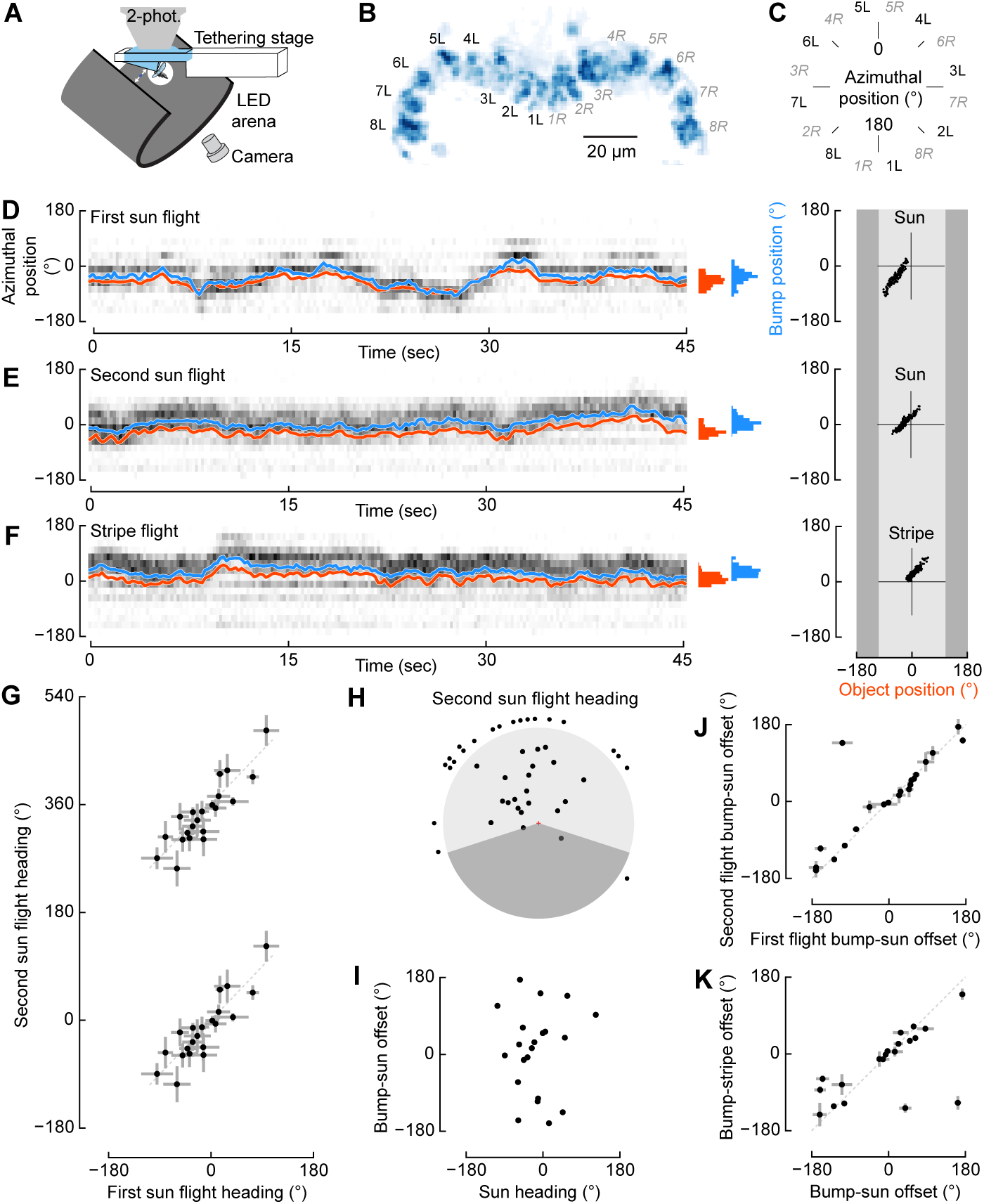
E-PG neuron activity correlates with both sun and stripe position. (A) Ca2+-imaging schematic. (B) Glomeruli assignment in protocerebral bridge based on a standard deviation of GCaMP6f fluorescence in E-PG terminals. (C) Continuous azimuthal representation of glomeruli in B. (D) GCaMP6f fluorescence (ΔF/F), represented as the unwrapped glomerular positions from C, during 45 seconds of a sun presentation (grey). Azimuthal position of E-PG activity bump (blue trace and probability distribution) and sun position (computed as in (25), red) co-vary. Regression of object position on the 216°-wide LED arena (light gray) against bump position for sequence plotted on left. (E) Similar to D, but for second sun presentation and (F) subsequent stripe presentation. (G) Heading during first sun trial plotted against heading in second sun trial with a minimum of 5 min inter-trial interval (N=20; plotted as in Figure 1E). (H) Polar representation of second sun bout headings, similar to Figure 1D. Shaded area not visible by fly. (I) E-PG bump-to-stimulus offset for each second sun flight bout. (J) Regression of the median bump-to-stimulus offset for first sun bout against the offset for the second sun bout. (K) Bump-to-stimulus offset for stripe regressed against the offset for the sun.

Based on well-established anatomy, we re-mapped the neural activity in the medial 16 glomeruli of the protocerebral bridge into the circular reference frame of azimuthal space (Fig. 2C, *27*) and computed a neural activity vector average, or bump position, for each image (similar to *28*; see Materials and Methods for details).

As in our flight arena experiments (Fig 1A), flies adopted arbitrary headings with respect to the sun stimulus (Fig. 2G, H), which they maintained over a 5-minute break (Fig 2G). By presenting sun and stripe stimuli to the same fly, we tested whether these two stimulus types are represented differently by the E-PG neurons. Bump position faithfully tracked the position of both the sun and stripe stimuli (Fig. 2D-F). Prior studies found that while the E-PG bump tracks the azimuthal position of a vertical stripe, it does so with an arbitrary azimuthal angular offset (*18*). We found an identical result with the sun stimulus; the bump rotated with changes in sun position, but with a bump-to-stimulus offset that varied from individual to individual. In addition, the bump-to-stimulus offset did not differ between the first and second sun presentation trials or between the sun and stripe presentation trails (Fig 2J, K). The offset was not correlated with the azimuthal angle at which individual flies tended to hold the sun (Fig. 2I). Together, these imaging results suggest that the representation of the sun and stripe in the E-PG neurons is similar despite the distinct behavioral responses to the stimuli, and that the bump-to-stimulus offset does not encode heading preference.

We next tested the causal contributions of E-PG neurons to sun navigation and stripe fixation, predicting that the highly variable headings adopted in sun navigation might require the instantaneous positional information provided by E-PG neurons. We took advantage of the sparse expression patterns of three different split-GAL4 lines (Fig. 3A) to selectively drive the inwardly rectifying potassium channel Kir2.1 (*29*). As a control, we crossed UAS-Kir2.1 to an engineered split-GAL4 line that was genetically identical to the experimental driver lines, but carried empty vectors of the two GAL4 domains in the two insertion sites (*30*). Driving Kir2.1 in three, separate split-GAL4 lines yielded flies that lost the ability to maintain the sun at arbitrary azimuthal positions, although they could fixate the sun and stripe frontally. To assess the degree to which this effect could have occurred by chance, we employed a bootstrapping approach similar to that used in our time gap experiments. We randomly selected 50 values from our control dataset 10,000 times, in each case calculating the circular variance of the subsampled population. We then determined the proportion of bootstrapped mean variances that had smaller values than the variance of the actual experimental data and concluded that the observed frontal distributions of our experimental groups were highly unlikely to have occurred by chance (SS00096: p=0.000; SS00408: p=0.000; SS00131: p=0.004). Thus, E-PG neuron activity appears necessary for menotaxis, i.e. maintaining the sun in arbitrary non-frontal positions. To our knowledge, this is the first behavioral deficit elicited via experimental manipulation of the compass cell network.

**Fig. 3.**
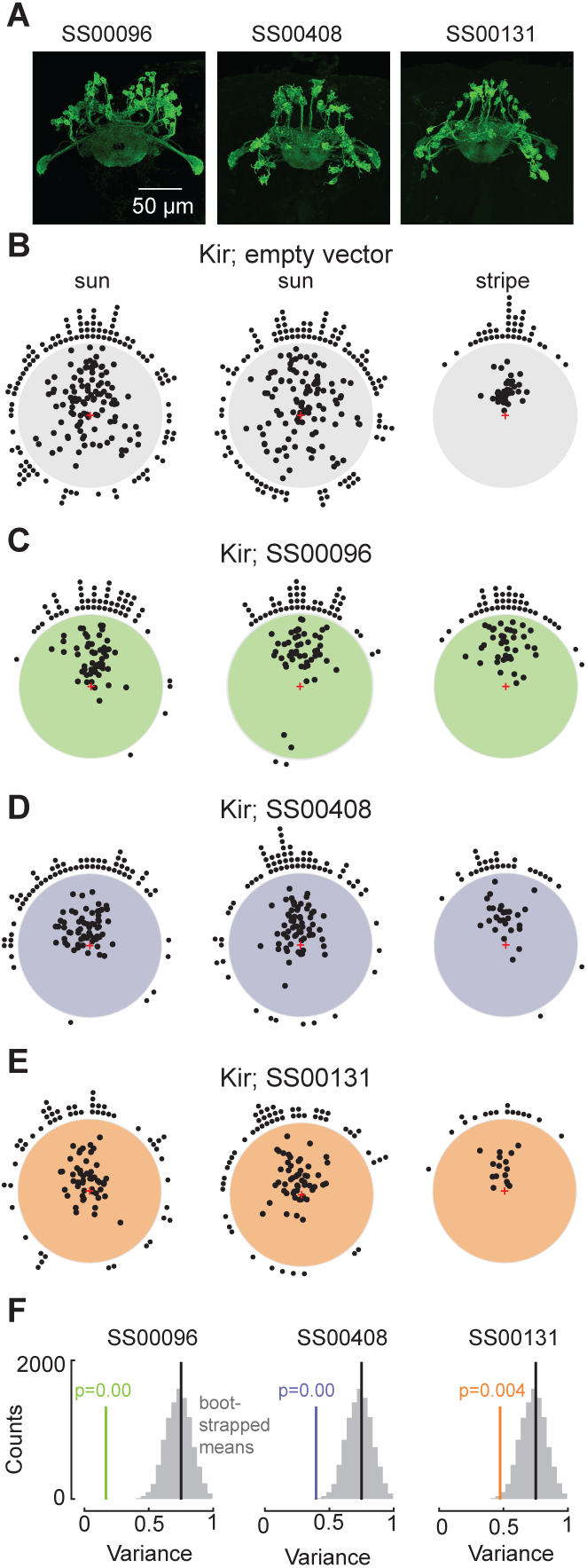
E-PG neuron activity is necessary for proportional navigation using sun. (A) Fluorescence labeling of GFP expressed in E-PG neurons in three experimental split-Gal4 lines, maximum intensity projections. (B) Sun menotaxis and stripe fixation in genetic controls (Kir; empty vector split-Gal4). (Left) First 5-min trial of sun fixation (n = 111 flies). (Center) Second 5-min trial of sun fixation (N = 108). (Right) Five minutes of stripe fixation (N = 38). (C) Results from the same experimental paradigm for Kir;SS00096 (N = 54, 49, 28). (D) Sun and stripe fixation for Kir;SS00408 (N = 64, 66, 28). (E) Headings from SS00131;Kir (N = 54, 54, 16). (F) Flies with silenced E-PG neurons have smaller variances than the genetic control. Distribution of 10,000 bootstrapped circular variances subsampled from the empty-vector control’s second sun trial, shown in gray (N = 50 each). Black lines depict the observed heading variance of the entire dataset (n = 108). Red lines indicate population heading variance of the second sun trial for each experimental group. P-value indicates the proportion of bootstrapped variances that are smaller than the observed variance for each experimental group.

In the absence of normal E-PG function, flies might directly orient toward the sun because they lack the ability to compare their instantaneous heading to a stored value of their directional preference. Such a loss of function in the compass network might unmask a simpler reflexive behavior – phototaxis – that does not require the elaborate circuitry of the central complex. Consistent with this hypothesis, stripe fixation was not different between control and experimental animals. This interpretation is compatible with a recent model that showed frontal object fixation could result from a simple circuit involving two asymmetric wide-field motion integrators, without the need for the central complex (*31*).

Our findings are consistent with an emerging model of a navigational circuit involving the central complex. E-PG cells have an excitatory relationship with another cell class in the central complex (protocerebral bridge-ellipsoid body-noduli or P-EN neurons), creating an angular velocity integrator that allows a fly to maintain its heading in the absence of visual landmarks (*19, 20*). Furthermore, the E-PG neurons are homologous to the CL1 neurons described in locusts (*32*), monarchs (*16*), dung beetles (*4*), and bees (*33*) and likely serve similar functions across taxa. In an anatomy-based model of path integration in bees, CL1 neurons are part of a columnar circuit that provides instantaneous heading information to an array of self-excitatory networks that also receive convergent optic flow information, thereby storing a memory of distance traveled in each direction (*33*). This information is then retrieved as an animal returns home by driving appropriate steering commands in other classes of central complex neurons. The putative memory cells suggested by this model, CPU4 cells, could be homologous to protocerebral bridge-fan shaped body-noduli (P-FN) neurons described for *Drosophila* (*27*). Furthermore, cells responsive to progressive optic flow are found throughout the central complex of flies, including neuropil in the fan-shaped body containing the P-FN cells (*34*). The authors of the recent path integration model suggest that the CPU4 network in bees might also function to store the desired heading during sun navigation (*33*). Although our results do not directly test this model, they are consistent with the role of CL1 neurons in providing heading direction to circuits that generate steering commands towards an arbitrary orientation whose memory is stored in the network of CPU4 (P-FN) neurons.

Stripe fixation and sun navigation behaviors may represent two different flight modes in *Drosophila*. Stripe fixation is thought to be a short-range behavioral reflex to orient towards near objects (*12*), which in free flight is quickly terminated by collision avoidance (*13*) or landing behaviors (*35*). In contrast, navigation using the sun is likely a component of long-distance dispersal behavior that could be used in conjunction with polarization vision (*8, 9*) either in a hierarchical (*4*) or integrative (*36*) manner.

Individuals could differ in where they lie on the continuum of long-range dispersal to local search, which could explain the inter-individual variation we observed in heading fidelity during sun orientation experiments. In general, dispersal is a condition-dependent behavior that is known to vary with hunger or other internal factors (*37*).

Given the architectural similarity of the central complex among species (*14*), the celestial compass we have identified in *Drosophila* is likely one module within a conserved behavioral toolkit (*13*) allowing orientation and flight over long distances by integrating skylight polarization, the position of the sun or moon, and other visual cues. An independent study has recently found that the E-PG compass neurons are also necessary in walking flies for maintaining arbitrary headings relative to a small bright object (*38*). The expanding array of genetic tools developed for flies as well as the rapid growth in our understanding of the neural circuitry involved in orientation during walking (*18–20*) and flight (*28*) make this a promising system for exploring such essential and highly conserved behaviors.

## Acknowledgments

We wish to thank Tanya Wolff and Gerry Rubin for generously providing us with the split-GAL4 lines SS00131 and SS00408 prior to the publication of their manuscript describing them. Crystal Liang and Aisling Murran provided valuable assistance with data collection. **Funding:** This work was funded by grants from the NSF (IOS 1547918), NIH (U19NS104655), and the Simons Foundation (71582123) to MHD, as well as an NIH NRSA postdoctoral fellowship (F32GM109777) to YMG. **Author contributions:** PTW and TLW were involved in early experimental design and analysis. YMG, KJL, IKR and MHD conceived of and conducted experiments. YMG characterized sun compass behavior (Fig. 1), IKR conducted functional imaging experiments (Fig. 2), and YMG and KJL performed genetic silencing experiments (Fig. 3). YMG, KJL, IKR and MHD wrote the paper. All authors contributed in editing the final manuscript. **Competing interests:** Authors declare no competing interests. **Data and materials availability:** All data will be made available on Dryad upon publication.

## Supplementary Materials

### Materials and Methods

#### Experimental animals

We conducted all experiments using 2-4 day old female *Drosophila melanogaster.* Our initial analysis of sun orientation behavior (Fig. 1) was conducted using flies from a wild-caught strain (‘top banana’) collected in Seattle, WA and maintained in the lab since September 2013. We reared flies in incubators on a 12 hour light: 12 hour dark cycle at 25°C on standard cornmeal fly food. For the functional imaging experiments (Fig. 2), we used flies heterozygous for w+;UAS-tdTomato;UAS-GCaMP6f (*39*) and the split-Gal4 line SS00096 (*28*). For silencing experiments (Fig. 3), we crossed a backcrossed version of UAS-Kir2.1 with SS00096, SS00131, and SS00408 (kindly provided by Tanya Wolff at Janelia Research Campus). The controls in our silencing experiments were the progeny of the UAS-Kir2.1 line and an engineered split-GAL4 line in which the two insertion sites carried empty vectors of the two GAL4 domains, but were otherwise genetically identical to the experimental driver lines (*30*). We generated flies for confocal imaging by crossing UAS-myr:GFP; UAS-red-Stinger with each of the split-GAL4 lines.

#### Fly tethering

For sun orientation (Fig. 1) and genetic silencing experiments (Fig. 3), we tethered flies under cold anesthesia and glued them to a tungsten wire (0.13 mm diameter) at the anterior dorsal portion of the scutum with UV-cured glue (Bondic Inc.). We also fixed the head of each fly to its thorax by applying an additional drop of glue. Flies were allowed to recover for at least 10 minutes prior to testing.

For functional imaging experiments (Fig. 2), we tethered each fly to a specially designed physiology stage (*40*) that permitted access to the posterior side of the fly’s head. We filled the holder with saline, and removed a section of cuticle overlying the region of the central complex. To improve imaging quality, we removed adipose bodies and trachea from the light path. Flies were continuously perfused with saline (*41*) which was actively regulated to a temperature of 21°C. We allowed flies a minimum of 20 minutes to recover from cold-anesthesia prior to imaging.

#### Flight arenas and stimulus presentation

For sun-orientation behavior (Fig. 1) and genetic silencing experiments (Fig. 3), we placed tethered flies in an LED flight simulator (*42*) (Fig. 1A). We displayed patterns on a circular arena of either 12 × 1 (Fig. 1) or 12 × 2 (Fig. 3) LED panels, with each panel consisting of an 8 × 8 array of individual pixels (Betlux #BL-M12A881PG-11, λ=525 nm). Each pixel subtended an angle of 2.8° at the center of the arena with a 0.93° gap between adjacent pixels. The panels were controlled using hardware and firmware (IORodeo.com) as described previously (*42*), with slight modifications in current sinking required to display a single bright pixel without generating bleed-through on other pixels in the same panel row. We placed the fly in the center of the arena at a body angle of ~60°, approximating the orientation during free flight (*43*). For wing tracking, flies were backlit with a collimated infrared source (850 nm, 900mW; Thorlabs Inc. #M850L3). We placed a 45° mirror below the fly and used a firewire camera (Basler A602f-2) or a Point Grey USB 3.0 camera (now FLIR, Blackfly 0.3MP monochrome camera, BFLY-U3-03S2M-CS) for image capture. Each camera was equipped with a Computar macro lens (MLM3X-MP) and IR-pass filter (Hoya B-46RM72-GB) to exclude extraneous light from the LED display.

To track the wing stroke envelope during flight, we used Kinefly, real-time machine-vision software developed in the lab (*44*). As in previous studies (*12*), we used the difference in wing beat amplitude (∆WBA) as a feedback signal by which the fly could control the angular velocity of the visual stimulus. The gain of this control relationship was set to 14.67, 5.88, or 4.75° sec^-1^ for each °∆WBA for sun orientation experiments (Fig. 1), functional imaging (Fig. 2), and genetic silencing (Fig. 3), respectively. We found that a lower feedback gain was required in our experiments with transgenic lines to generate stable orientation behavior to both sun and stripe stimuli.

For functional imaging experiments, we presented visual stimuli using a 12 × 4 panel (96 × 32 pixel) arena, which covered 216° of azimuth with a resolution of ~2.25°. To reduce light pollution from the LED arena into the photomultiplier tubes of the 2-photon microscope, we shifted the spectral peak of the visual stimuli from 470 nm to 450 nm by placing two transmission filters in front of the LEDs (Roscolux no. 59 Indigo and no. 39 Skelton Exotic Sangria). We tracked wing stroke angles using Kinefly and presented stimuli in closed-loop as described above, except that we illuminated the wings using four horizontal fiber-optic IR light sources (Thorlabs Inc. #M850F2) distributed in a ~90° arc behind the fly.

For data presented in Figs. 1 and 3, a single pixel served as our ersatz sun. At the plane of the fly, a single pixel subtends a maximum angle of 2.8°. However, because the fly was placed ~30° below the plane of the illuminated pixel, the simulated sun subtended a maximum angle of ~2.3° at the fly’s retina, which is larger than the sun’s angular diameter ( ~0.5°), but smaller than the inter-ommatidial acceptance angle of ~5° (*45*). For sun orientation experiments (Fig. 1), we conducted all trials in a 12 × 1 panel (96 × 8 pixel) arena. For stripe fixation, we presented a 4 pixel-wide dark stripe (15° wide × 30° high) on a bright background.

To determine the visual contrast flies experienced during our experiments, we measured the normalized difference between the lightest and darkest parts of the display (Michelson contrast). We placed a small metal tube covered in black electrical tape over a power sensor (Thorlabs S170C, PM100D) to reduce reflections and approximate the acceptance angle of an ommatidium (~5°). We held the sensor at the center of the arena and directed it toward a sun or stripe to measure the stimulus light level and then moved the stimulus 45° in azimuth to measure the background light level. The Michelson contrast for all sun stimulus experiments was 0.99 and stripe contrast was 0.75 and 0.74 for the data presented in Fig. 1 and Fig. 3 respectively.

The behavior of flies from the control line (UAS-Kir; split-GAL4-empty-vector) was generally similar to wild type flies; however, they tended to perform poorly in the stripe-fixation paradigm, as indicated by relatively low vector strength (Fig. 3B) and a smaller proportion of flies that completed the trial without stopping. Given that flies’ azimuthal control of a stripe stimulus improves as a function of increasing stripe height (*24*), we doubled the height of the visual display to a stripe of ~58° (12 × 2 panels, 96 × 16 pixels) for our genetic silencing experiments (Fig. 3). We noted that reflections generated by a single bright pixel on the faceted inner surface of the arena generated a faint dark stripe on the column of panels on which the sun stimulus was displayed. To guard against the possibility that the fly would orient to this reflection feature, we fabricated cylindrical inserts of black velvet that obscured the surface of the display except for a narrow slit (9 mm × 360°) that contained the LED row in which the sun stimulus was displayed. The insert could be quickly removed without disturbing the fly for trials using a stripe stimulus. To facilitate the collection of large sample sizes for the genetic silencing experiments, we constructed two identical arenas, which we operated in parallel.

In the functional imaging experiments (Fig. 2) we compensated for a larger arena and dimmer LEDs by using a 3.6° × 3.6° spot (2 × 2 pixels) as our sun stimulus, with a Michelson contrast of 0.92. The stripe stimulus consisted of a 12.6° wide × 64° high vertical bright stripe presented on a dark background, with a Michelson contrast of 0.93. Using a bright stripe on a dark background was necessary in order not to saturate the PMTs. Flies exhibit less robust fixation under these conditions (*42*) but nevertheless performed the behavior, allowing us to compare sun- and stripe-fixation during functional imaging.

#### Sun orientation and time gap experiments

To test the persistence of flight headings, we presented flies with the sun stimulus in closed loop, provided a rest period between flights for either 5 minutes, 1 hour, 2 hours, or 6 hours, and then tested flies in a second bout with a sun stimulus. Before and after the sun stimulus trials, we presented flies with a stripe for 1 minute. For 5-minute inter-trial intervals, we left the fly in the arena and stopped flight by presenting a small piece of paper. To prevent dehydration during longer inter-trial intervals (1, 2 and 6 hours), we removed the fly from the arena and placed it on a small foam ball floating in a microcentrifuge tube filled with water. Following this rest period, we returned the fly to the arena and the second flight was initiated by providing a small puff of air. We discarded trials in which any fly stopped flying more than once during any of the stripe or sun presentations.

#### 2-photon functional imaging

We imaged at an excitation wavelength of 930 nm using a galvanometric scan mirror-based two-photon microscope (Thorlabs, Inc., Newton, NJ, USA) equipped with a Nikon CFI Plan Fluorite objective water-immersion lens (10x mag., 0.3 N.A., 3.5 mm W.D.). With the addition of a piezo-ceramic linear objective drive (P-726, Physik Instrumente GmbH and Co. KG, Karlsruhe, Germany) we imaged two x-y planes separated by 25 μm along the z axis. Within the resulting volume we recorded tdTomato and GCaMP6f fluorescence in those glomeruli of the protocerebral bridge (PB) that contain terminals of E-PG neurons. We scanned in a boustrophedon pattern from ventral to dorsal to align the piezo drive descent during each plane scan with the anatomical inclination of the PB, maximizing the volumetric capture of the target glomeruli. We acquired the 142 × 71 μm images with 128 × 64 pixel resolution at 13.1 Hz. The 2-plane scan with one fly-back frame resulted in a 4.36 Hz volumetric scan rate. To correct for motion in the x-y plane, we registered both channels for each frame by finding the peak of the cross correlation between each tdTomato image and the trial-averaged image (*46*). Subsequently, we collapsed the two planes with a maximum z-projection. Based on known anatomy, we manually assigned a region of interest (ROI) to each PB glomerulus with E-PG neuron innervation. For each volumetric frame, we computed fluorescence (F_t_) of the GCaMP6f signal by subtracting the mean of the background pixels from the mean of the ROI pixels for each glomerulus. The background was defined as the 10% dimmest pixels across the entire z-projected image for each fly. We normalized the fluorescence in the ROI of each glomerulus to its baseline fluorescence (F_0_) as follows: ΔF/F = (F_t_ – F_0_)/F_0_ and defined F0 as the mean of the 10% lowest GCaMP6f fluorescence in the ROI of each glomerulus. Under closed-loop conditions, we presented each fly with a sun stimulus twice for five minutes, separated by a minimum of 5 minutes. A 2-minute presentation of the stripe stimulus followed the second sun stimulus trial.

#### Functional silencing of E-PG neurons in sun navigation behavior

We tested all control and experimental flies with a paradigm consisting of 5 minutes of sun stimulus presentation, a 5-minute break, 5 minutes of sun presentation, and 5 minutes of stripe presentation. We discarded trials in which flies stopped more than twice per stimulus presentation.

#### Immunohistochemistry of split-GAL4 lines

We dissected and stained brains of flies expressing UAS-myr:GFP, UAS-redStinger and each of the three split-GAL4 lines (SS00096, SS00131, SS00408) using modifications to standard laboratory immunohistochemistry protocols (*44*). We dissected brains in 4% formaldehyde fixative. After a 20-30 minute fixation, we washed tissue 2 × 20 minutes in phosphate buffered saline (PBS) followed by a permeabilization step of 2 × 20 minute washes in phosphate buffered saline with 0.5% Triton-X (PBST). We incubated tissue with primary antibodies anti-GFP AlexaFluor^™^ 488 conjugate (1:1000 concentration, Invitrogen # A21311) and anti-nc82 to label neuropil (1:10 concentration, Developmental Studies Hybridoma Bank, AB 2314866) in 5% normal goat serum in PBST overnight on a nutator at 4°C. The following day, we washed with PBST 3 × 20 min and incubated with a secondary antibody to anti-nc82 (AlexaFluor^™^ 633, 1:250 concentration, Invitrogen # A21050) overnight at 4 °C. Brains were washed 3 × 20 min with PBST and 2 × 20 min with PBS the following day. We dehydrated brains through an ethanol series (30%, 50%, 70%, 90%, 100%, 100%, each for 10 min), cleared tissue with xylene (2 × 10 min) and mounted in DPX (*47*). Using a Leica SP8, we imaged brains under a 63x objective (Leica #506350, 1.4 N.A.). Maximium intensity projections were generated in Fiji (*48, 49*).

#### Quantification and statistical analysis

We processed and analyzed all data in Python 2.7 and Matplotlib (*50*). Before making pairwise comparisons of mean heading direction in separate flights (as in Fig 1E, F), we excluded trials with a vector strength under 0.2 (36.2% of all trials). Mean headings for flights with very low vector strength are not meaningful, as this indicates that the fly did not select a heading during the trial. However, including all data did not qualitatively change the relationship between first and second flights.

To assess whether heading fidelity was maintained over time gaps, we bootstrapped random pairings of first and second sun presentation trials 10,000 times. We compared the distribution of the mean absolute value of heading difference between the flights for these simulated data sets to the mean absolute value of heading difference of the observed data. We calculated the p-value as the proportion of simulated data sets that had a mean heading difference smaller than that of the observed data. We conducted a similar analysis for the results of our behavioral genetics experiments. In that case we bootstrapped subsamples (N=50) of our control dataset with replacement 10,000 times and calculated the circular variance of each dataset. As above, we then reported the proportion of bootstrapped data sets with a smaller variance than each experimental group. We selected a resample size of 50 as this approximated the sample size of our datasets (N=49, 54, 64). A systematic analysis of p-values showed that they decreased asymptotically to a constant level at resample sizes greater than 20.

#### Data and software availability

Data will be uploaded to Dryad.

### Movie S1

#### E-PG activity correlates with azimuthal position of sun and stripe visual objects under open-loop flight conditions

All panels are time-synchronized and sampled at the two-photon volumetric imaging rate. Upper panel: Two-photon Ca^2+^ imaging in the protocerebral bridge of a flying fruit fly. The fly is presented with sun and stripe stimuli that move in azimuth at a constant speed (open loop). Upper 10% GCaMP6f fluorescence (ΔF/F) in green, lower 90% ΔF/F in grey; Gaussian filtered. Middle panel: wing tracking (red) in the machine vision camera view of the tethered fly. This ventral view is flipped to represent the anatomical right wing on the right. Stimulus position (blue) and E-PG neuron activity bump position (green) represented in azimuthal space (not to scale). Lower panels: ΔF/F during 20 seconds of sun and 20 seconds of stripe stimuli presentations. White vertical stripe indicates current frame. Azimuthal position of E-PG activity bump (blue dots and probability distribution) and stimulus position (computed as in (*1*), red) co-vary. Visual stimuli are only presented on the centered 216°-wide LED arena.

### Movie S2

#### E-PG activity correlates with azimuthal position of sun and stripe visual objects under closed-loop flight conditions

Identical representation to Movie S1, but with stimuli presented under closed loop conditions (see Materials and Methods section for details).

